# Proteomic Stratification of Prognosis and Treatment Options for Small Cell Lung Cancer

**DOI:** 10.1101/2023.10.22.563494

**Authors:** Zitian Huo, Yaqi Duan, Dongdong Zhan, Xizhen Xu, Nairen Zheng, Jing Cai, Ruifang Sun, Jianping Wang, Fang Cheng, Zhan Gao, Caixia Xu, Wanlin Liu, Yuting Dong, Sailong Ma, Qian Zhang, Yiyun Zheng, Liping Lou, Dong Kuang, Qian Chu, Jun Qin, Guoping Wang, Yi Wang

## Abstract

Small cell lung cancer (SCLC) is a highly malignant and heterogeneous cancer with limited therapeutic options and prognosis prediction models. Here, we analyzed formalin-fixed, paraffin-embedded (FFPE) samples of surgical resections by proteomic profiling, and stratified SCLC into three proteomic subtypes (S-I, S-II, and S-III) with distinct clinical outcomes and chemotherapy responses. The proteomic subtyping was an independent prognostic factor and performed better than current TNM or Veterans Administration Lung Study Group (VALG) staging methods. The subtyping results could be further validated using FFPE biopsy samples from an independent center, extending the analysis to both surgical and biopsy samples. The signatures of the S-II subtype in particular suggest potential benefits from immunotherapy. Differentially overexpressed proteins in S-III, the worst prognostic subtype, allowed us to nominate potential therapeutic targets, indicating that patient selection may bring new hope for previously failed clinical trials. Finally, analysis of an independent cohort of SCLC patients who had received immunotherapy validated the prediction that the S-II patients had better Progression Free Survival (PFS) and Overall Survival (OS) after first-line immunotherapy. Collectively, our study provides the rationale for future clinical investigations to validate the current findings for more accurate prognosis prediction and precise treatments.

## Introduction

Small cell lung cancer (SCLC) is an exceptionally aggressive lung neuroendocrine neoplasm characterized by rapid tumor growth, early metastasis, and acquired chemo-resistance. Comprehensive whole-exome and whole-genome analyses have identified inactivation of *TP53* and *Rb1* as the predominant genetic alterations in SCLC, occurring in > 98% of the patients [1,2]. Genomic analysis of 51 SCLC cases also found genetic alterations in the PI3K/AKT/mTOR pathway in 36% of the samples, with mutations in *PIK3CA* (6%), *PTEN* (4%), *AKT2* (9%), *AKT3* (4%), *RICTOR* (9%), and *mTOR* (4%) [3]. However, effective stratification markers and treatment targets for SCLC remain limited [4]. As a result, the overall survival (OS) of SCLC patients has seen no significant improvement despite numerous clinical trials of different chemotherapy schemes and biological agents over the past decades [5]. Moreover, a lack of biomarkers that help predict efficacy has also impeded SCLC patients from reaping significant benefits from immunotherapy [6]. Currently, the 5-year survival rate is approximately 20-25% for limited-stage SCLC (LS-SCLC) and barely 1–2% for extensive-stage SCLC (ES-SCLC), making SCLC one of the deadliest cancers.

Early studies based on cell line morphologies classified SCLC into classic subtypes that expressed higher neuroendocrine (NE) markers and variant subtypes that showed low or an absence of neuroendocrine features [7,8,9,10]. These characteristics were further observed in clinical samples [11,12]. Subsequently, SCLC was classified according to the expression of neuroendocrine transcription factors ASCL1 and/or NEUROD1 [13]. Additionally, POU2F3 expression was used to define a non-NE, tuft-cell variant of SCLC [14]. And although YAP1 was proposed as a potential subtype marker for the remaining unclassified SCLC cases [3], it has yet to be confirmed [15]. Recently, re-analyzing previously published transcriptomic data classified SCLC into four subtypes [16]; in addition to the previously identified subtypes with the ASCL1 (SCLC-A), NeuroD1 (SCLC-N), and POU2F3 (SCLC-P) signatures, a new subtype (SCLC-I) characterized by the expression of inflammation gene signatures was uncovered.

In IMpower133, a global phase I/III, double-blind, randomized, placebo-controlled trial, atezolizumab (anti–PD-L1) was added to carboplatin + etoposide for ES-SCLC. SCLC-I patients were reported to experience the greatest benefit from this combined immuno- and chemotherapy. In the randomized, controlled, open-label, phase 3 CASPIAN trial, first-line durvalumab plus platinum-etoposide also significantly improved overall survival in patients with ES-SCLC versus a clinically relevant control group [17]. Additionally, cisplatin treatment of SCLC-A patient-derived xenografts induced intratumoral shifts toward SCLC-I [16]. These analyses suggest that the SCLC-I subtype might be the right candidate for immunotherapy. Therapeutic vulnerabilities were also identified for each subtype, including to inhibitors of PARP (SCLC-P), Aurora kinases (SCLC-N), or BCL-2 (SCLC-A) [16]. More recently, single-cell transcriptome sequencing (scRNA-seq) and imaging techniques have further revealed the heterogeneity and tumor microenvironment (TME) of SCLC [18]. Monocytes/macrophages appear to play a profibrotic and immunosuppressive role in SCLC TME. SCLC-N showed less immune infiltrate and greater T cell dysfunction than SCLC-A. More importantly, most SCLC cases share a small PLCG2-high subpopulation, which is linked to metastasis and poor prognosis.

While subtyping based on genomic and transcriptomic analyses has greatly improved our understanding of SCLC, the resulting subtypes correlated poorly with clinical outcomes. In contrast, subtyping with proteomics had revealed its exceptional clinical potential for more accurate predication of prognosis, chemo-sensitivity, and treatment targets for myriad cancers including stomach [19,20], liver [21,22], ovarian carcinoma [23], colorectal cancer [24], and NSCLC [25,26,27,28,29]. Here, we report a proteomic subtyping model derived from a discovery dataset containing 75 surgically resected formalin-fixed, paraffin-embedded (FFPE) samples. The model was then validated in an independent cohort of 52 FFPE biopsy samples. The subtypes based on our proteomic model correlated well with clinical information including OS, and also allowed subtype-specific nomination of drug targets in SCLC. In addition, analysis of another 52 samples from patients who received immunotherapy allowed us to validate the finding that one particular proteimic subtype is an immune responsive subtype.

## Results

### Limited predictive power of SCLC staging and classification systems

In an effort to develop a more reliable SCLC subtyping system, we first analyzed the 75 surgically resected formalin-fixed, paraffin-embedded (FFPE) SCLC samples as shown in **Figure 1**A. Key clinical characteristics of the patients in the discovery cohort are presented in Figure 1B, with their detailed clinical and pathological data provided in Table S1. Of the 75 cases, 62 (82.7%) were LS-SCLC and 13 (17.3%) were ES-SCLC according to the Veterans Administration Lung Study Group (VALG) definition. Based on the tumor-node-metastasis (TNM) staging system, 13 (17.3%), 19 (25.3%), 28 (37.3%) and 13 (17.3%) cases were categorized as stage I, II, III, IV, respectively. At the end of follow-up, 31 (41.3%) patients survived, with a median overall survival (OS) time of 2.78 ± 0.25 years. The statistically significant univariate prognostic factors included age (Log-rank *P* value = 0.041), gender (Log-rank *P* value = 0.0085), and lymph node metastasis (LNM (Log-rank *P* value = 0.041) (Figure S1).

**Figure 1.**
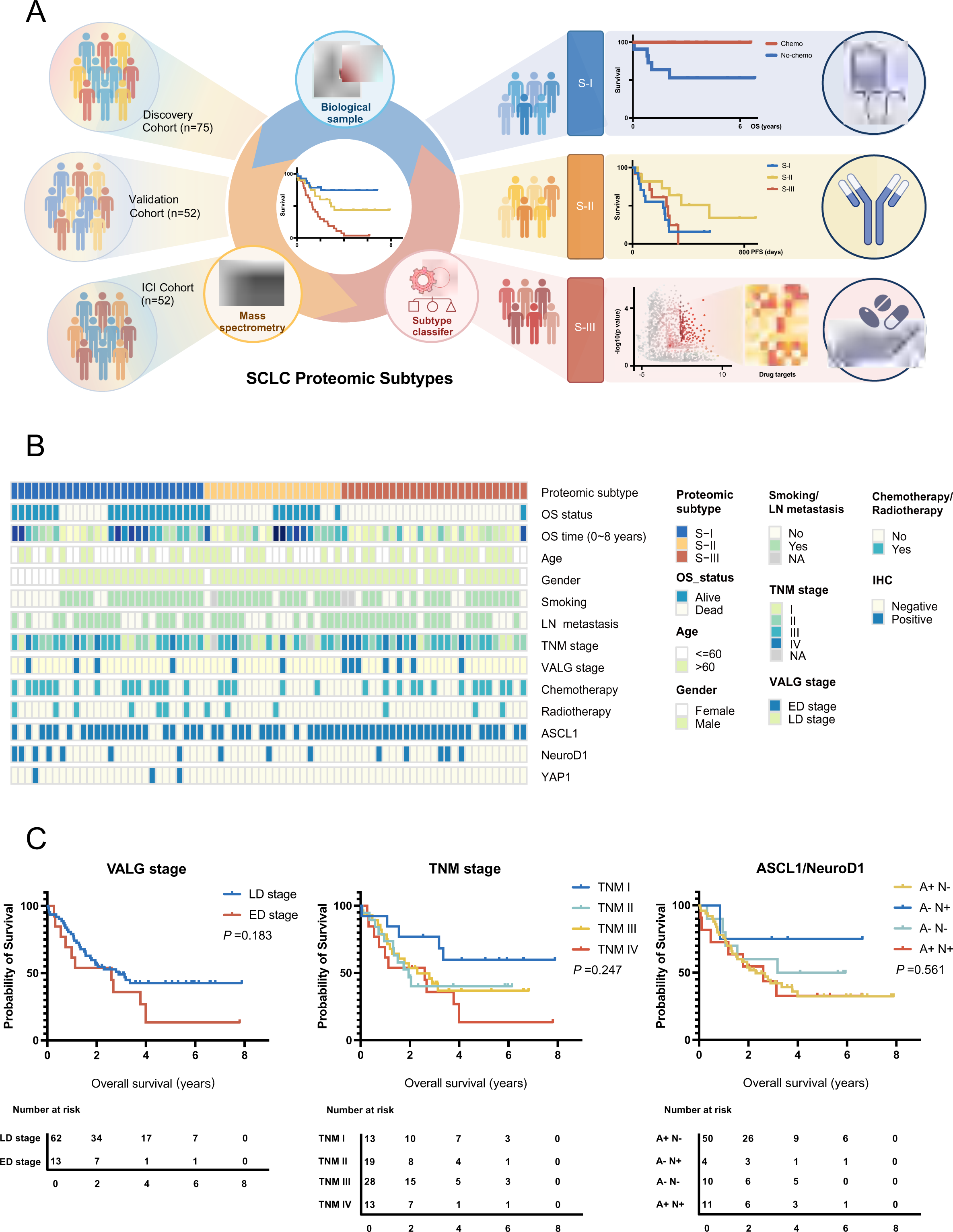
Study design and key clinical characteristics of the SCLC patients from the discovery cohort. **A.** A flowchart of our study scheme. **B.** clinical characteristics of the patients from the discovery cohort. **C.**The Kaplan-Meier plot shows the limited predictive value on patient overall survival of the VALG, TNM, and ASCL1/NeuroD1 staging and classification methods. VALG, Veterans Administration Lung Study Group;

When the predictive value of the current staging systems (including VALG and TNM) were examined against actual prognosis of SCLC patients, their limitations became clear (Figure 1C). In fact, only TNM I patients had significantly better OS than those of TNM IV (Log-rank *P*-value = 0.04) (Figure 1C). Immunohistochemistry (IHC) was used to examine the expression of ASCL1, NeuroD1, and YAP1 in the samples, and positive rates of 81.3% (61/75), 20% (15/75), and 4% (3/75), were obtained respectively. Of all cases, 11 were double-positive for ASCL1 and NeuroD1 (Figure S2 & Table S1). However, classification based on ASCL1/NEUROD1 expression did not correlate with OS differences in patients either (Figure 1C), although the NeuroD1^+^/ASCL1^-^ subtype showed better but not statistically significant prognosis (Log-rank test *P*=0.56).

### Proteomics stratifies SCLC into subtypes that correlate with clinical outcomes

Since the current SCLC staging and classification systems failed to make a satisfactory prediction, we carried out a proteomic study using these 75 samples by label-free quantitative mass spectrometry (Figure 1A). We detected a total of 6,108 gene products of high confidence from 75 samples, with 2957 proteins identified in more than 50% samples. The protein identities and relative abundances of each case, designated as fraction of total (FOT) [30], were provided in Table S2. To develop a robust classification model, we first selected the top 1,100 most abundant proteins detected from each sample, which yielded a dataset of 3,460 proteins; then the 445 proteins that were detected in at least 7 samples (>10%) with the coefficient of variations (CV) greater than 1.9 were used for non-negative matrix factorization (NMF) consensus clustering (Table S3). NMF clustering yielded three subgroups, namely S-I (n=28, 37%), S-II (n=20, 27%), and S-III (n=27, 36%) with the maximum average silhouette of 0.83 (**Figure 2**A). Importantly, these proteomic subtypes correlated well with overall survival prognosis. Patients in S-I had the best overall survival with a 5-year OS probability of 75%, whereas S-III had the worst survival with only 3.7% of 5-year OS (Log-rank *P*-value < 0.001) (Figure 2B). A multivariate Cox analysis confirmed that the proteomic subtype was an independent prognostic factor (S-I vs S-III, HR = 4.73, 95% CI, 1.81-12.4, Cox *P*-value = 0.002) after adjusting for TNM stage, VALG stage, and other covariates including chemotherapy, age, gender, LN metastasis status, and smoking history (Figure S3). These data indicated that proteomics-based subtyping is superior in overall survival prognosis.

**Figure 2.**
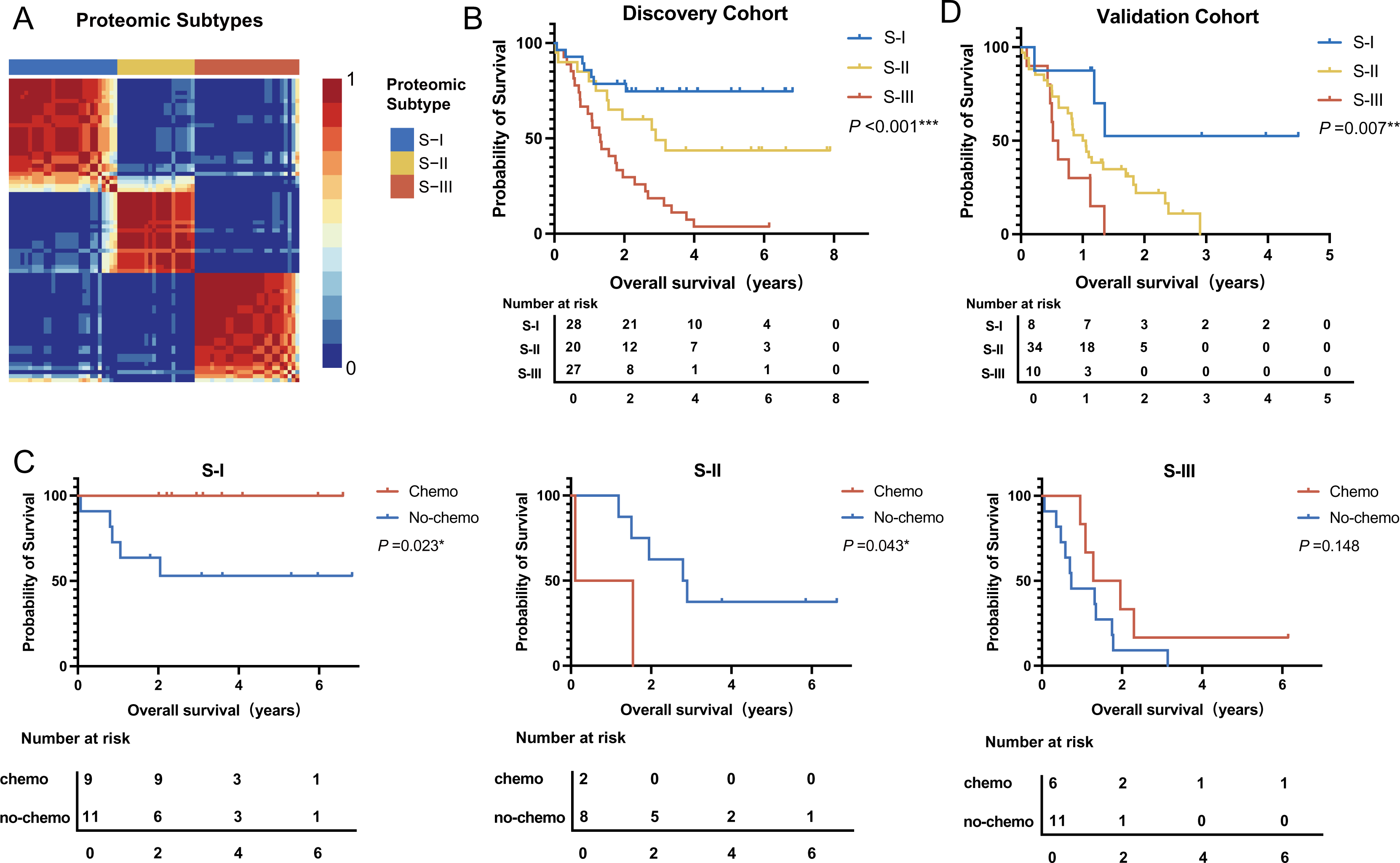
Proteomics stratifies SCLC into subtypes that correlate with clinical outcomes. **A.** In the discovery cohort, NMF clustering yielded three subgroups in SCLC. **B.** The three subtypes based on proteomic profiling were associated with different clinical outcomes. **C.** In the validation cohort, biopsy samples were obtained from the First Affiliated Hospital of Henan University. The predictive classifier model shows the prognosis trend similar to the discovery cohort. **D.** The three proteomic subtypes exhibited different responses to chemotherapy. S-I, Log-rank *P*-value = 0.023. S-II, Log-rank *P*-value = 0.042. S-III, Log-rank *P*-value = 0.15.

### The proteomic subtypes show different benefit of chemotherapy on prognosis

The platinum agent (cisplatin or carboplatin) and etoposide-based combination chemotherapy is the standard care for SCLC patients after surgery, although the percentage of patients who actually benefit from such care is quite small [31]. In our dataset, approximately half of the patients (33, 44%) underwent chemotherapy. Considering the impact of TNM I and IV on prognosis mentioned earlier, we included only TNM II and III patients for chemosensitivity analysis. As shown in Figure 2C, chemotherapy exhibited different impact on prognosis for the three proteomic subtypes. The S-I subtype benefited most significantly from chemotherapy (Log-rank *P*-value = 0.023), with all patients who received chemotherapy were still alive by the end of the study. The S-II subtype appeared to have worse prognosis (Log-rank *P*-value = 0.042). The S-III subtype had the worst prognosis and were chemo-insensitive (Log-rank *P*-value = 0.15). In summary, our proteomic SCLC subtyping could stratify patients into 3 distinct subtypes that are more clinically relevant. The S-I subtype was chemo-sensitive with the best OS. S-II had medium OS where chemotherapy might be detrimental. S-III had the poorest prognosis and was insensitive to chemotherapy, suggesting that S-III patients have the greatest need for new therapies.

To facilitate the validation of the subtyping in the independent external dataset, we developed a random forest (RF) classifier with the discovery set. The top 500 most abundant proteins detected in each sample were aggregated for differential expression analysis. The resulted 58 signature proteins (Fold change > 1.5, adjusted t-test P-value < 0.05) with high identification frequencies (detected in more than 25% of the samples) were used as input (predictor variables) (Table S4). We trained a random forest (RF) classifier with a 10-fold cross-validation in the discovery dataset. Based on the 58 features, the predictive classifier model yielded a 90.8% accuracy for the discovery dataset. We collected 52 SCLC biopsy samples from another center (the First Affiliated Hospital of Henan University) as an independent validation dataset. The corresponding clinical information of these samples is summarized in Table S5. This subset included 12 female and 40 male patients, with an overall survival time ranging from 0.019 to 4.493 years. The two-year survival rate was 8/52, or 15.4% and the three-year survival rate was 2/52, or 3.8%. Proteomic analysis were performed on the 52 biopsy samples. The results were listed in Table S6. When the RF classifier was applied to the validation dataset, the predicted S-I, S-II, and S-III subtypes contained 8, 34, and 10 cases, respectively. The prognosis trend was consistent with that of the discovery cohort, with one-year survival rates of 87.5%, 52.9%, and 30.0% and two-year survival rates of 37.5%, 14.7%, and 0% for S-I, S-II, and S-III, respectively. Kaplan-Meier analysis illustrated that the overall survival varied significantly among the three subtypes (Log-rank test *P* = 0.007) (Figure 2D). Thus, the proteomic subtyping model derived from surgical samples could be validated with biopsy samples from an independent center.

### Differential signature proteins and enriched biological processes in proteomic SCLC subtypes

To investigate the proteomic features of SCLC subtypes, we selected significantly altered proteins by comparing their expression in each subtype to the other subtypes (Wilcox test *P* < 0.05 and fold change > 3 for S-I and II or fold change > 10 for S-III). As a result, 206, 162, and 126 subtype-specific proteins for S-I, S-II, and S-III respectively were designated as significantly altered proteins. (**Figure 3**A, Table S7). Functional enrichment using Metascape showed that the most distinct subtype was S-II, which was significantly enriched in extracellular matrix, interferon gamma signaling and neutrophil degranulation (adjusted *P* values were between 10^-8^-10^-17^) (Figure 3B). In the regulation of immune functions, Major histocompatibility complex class I (MHC-I) molecules play an important role in cell-mediated immunity by presenting tumor antigens to CD8+ T cells and enabling cytotoxic T cells to recognize and eliminate tumor cells. Antigen presenting cells (APCs) such as B cells, dendritic cells (DCs), and monocytes/macrophages express major histocompatibility complex class II molecules (MHC II) and present antigenic peptides to CD4+ helper T cells. Here, we found that the MHC-I (including HLA-A, HLA-B, and HLA-C) but not the MHC-II (including HLA-DQA1, HLA-DQB1, HLA-DRA, HLA-DRB1, HLA-DRB5, HLA-F and HLA-E) molecules were expressed at higher levels in S-II than in the other two subtypes (Figure 3C & D). Moreover, TAP1, TAP2, and TAPBP were also highly overexpressed in S-II.

**Figure 3.**
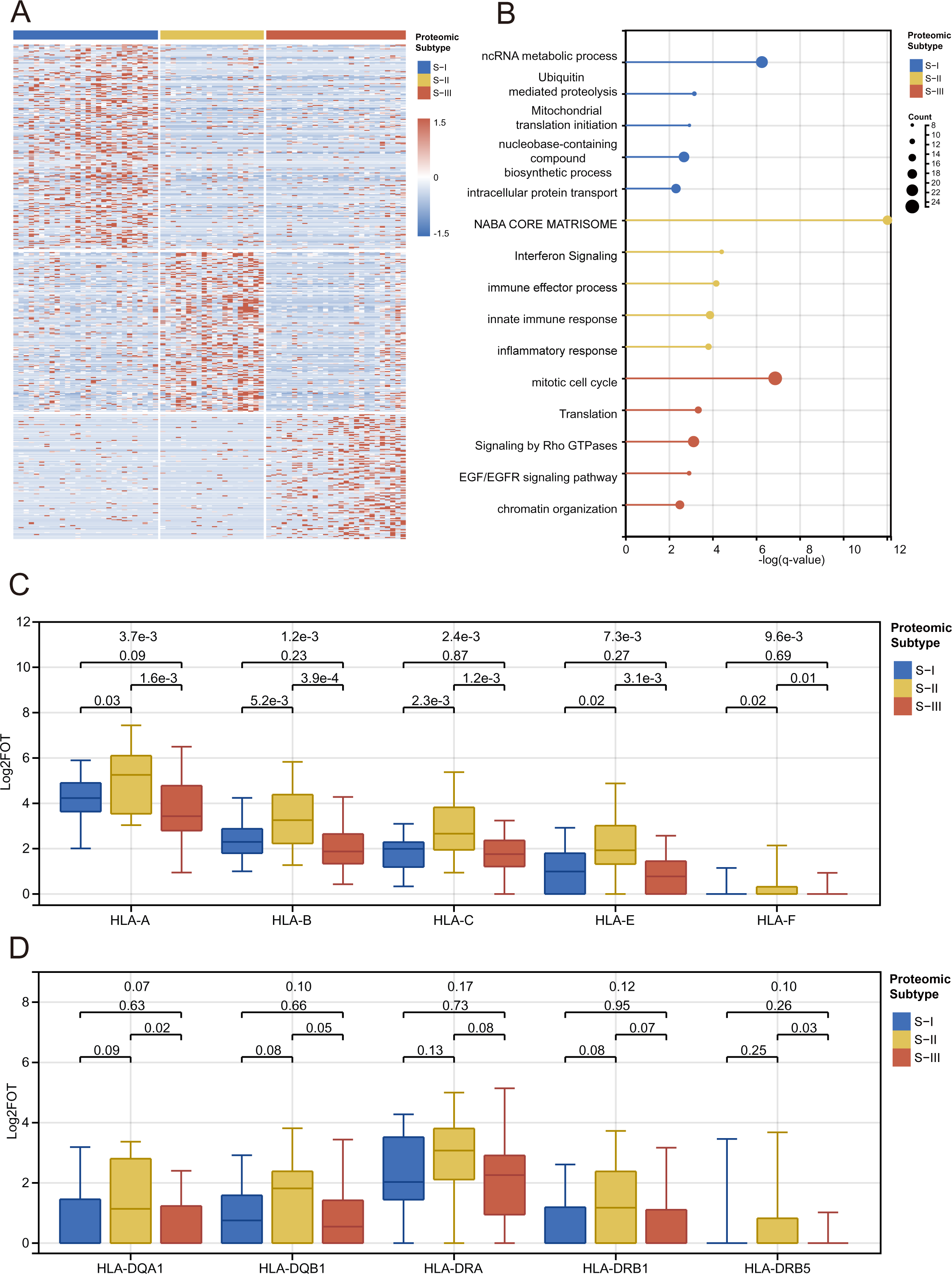
Differential signature proteins and enriched biological processes in proteomic SCLC subtypes. **A.** Heatmap of signature proteins in each subtype. **B.** Metascape functional enrichment analysis identified different biological processes or pathways activated in the three subtypes. **C-D.** the MHC-I(**C**) rather than MHC-II (**D**) molecules were highly expressed in S-II compared with S-I and S-III.

TAP1 and TAP2 are members of the superfamily of ATP-binding cassette (ABC) transporters that shuttle various molecules across extra- and intra-cellular membranes. The ATP-loaded TAP1-TAP2 complex mediates unidirectional translocation of peptide antigens from the cytosol to endoplasmic reticulum (ER) for their loading onto MHC class I (MHC-I) molecules [32]. TAPBP is a transmembrane glycoprotein that mediates interactions between newly assembled MHC-I molecules and TAP proteins, and is required for the transport of antigenic peptides across ER membrane[33]. Thus, the S-II subtype is characterized by an enrichment of factors involved in antigen presentation.

The S-I subtype appeared to be marginally enriched in proteins functioning in transport, membrane trafficking, and RNA metabolic processes (adjusted *P* values of 10^-2^-10^-4^). Detailed analysis identified ubiquitin-mediated proteolysis in cell cycle control, including the anaphase promoting complex (ANAPC1, ANAPC13, and ANAPC4) and COPS9 signalosome components (COPS3 and COPS7B) as well as NEDD8. DNA repair and DNA replication proteins, including MSH3, LIG1, POLE3, RFC3, ORC2, and ATM, were also enriched (Table S7). After adjusting for *P* values, Metascape analysis could not identify significantly enriched biological processes for S-III. Manual inspection showed that proteins in chromatin organization/transcription regulation as well as various enzymes were enriched (Table S7). Among them were proteins mediating chromatin assembly and telomere maintenance (CHAF1A, TERF2, and TOX4), transcription coactivators (e.g., ATAD2, ZNF516, PHF6, PHF8, MED6, TAF5, and TAF) and corepressors (e.g., TBL1X, TBL1Y, CHD8, ATXN2, and SPEN). It appears that mis-regulation in chromatin structure and transcription may profoundly impact an array of biological processes, which the Metascape algorithm might have failed to identify.

### The S-II patients had the best immunotherapy responses in SCLC

Our analysis above suggested that S-II was an inflamed SCLC subtype in nature, which may benefit from immunotherapy. To test this hypothesis, we collected another 52 real-world FFPE biopsy or surgery samples from ES-SCLC patients who received Immune checkpoint inhibitors (ICIs) (including Sintilimab, Toripalimab, Durvalumab, Camrelizumab, Tislelizumab, and Atezolizumab). Among them, 37 were treated with combined immunotherapy with chemotherapy as a first-line treatment, and 49 were from biopsy (**Figure 4**A & Table S8). We classified these samples using the model derived from the discovery dataset (Table S9) into S-I (23 patients), S-II (13 patients), and S-III (16 patients). We specified an Immune checkpoint inhibitors PFS (ICI-PFS) as the duration between the time when the first immunotherapy was applied, and Progressive Disease (PD) was determined by clinical standards. As shown in Figure 4B&C, the ICI-PFS of the S-II patients with first-line ICIs rather than > first line ICIs was significantly longer than those of S-III (Log-rank test *P* = 0.04), and S-I (Log-rank test *P* = 0.08). Moreover, when the K-M plots of the S-II patients in the immunotherapy cohort, the discovery cohort (WH-S-II, no immunotherapy) (Log-rank test *P* = 0.03) and the validation cohort (HN-S-II, no immunotheray) (Log-rank test *P* < 0.0001) (Figure 4D) were compared, the S-II patients achieved better OS when trated with immunotherapy. Together, these analyses demonstrated the values of stratifying patients for immunotherapy, and suggested that immunotherapy could significantly improve the OS of SCLC patients, particularly for those in the S-II subtype.

**Figure 4.**
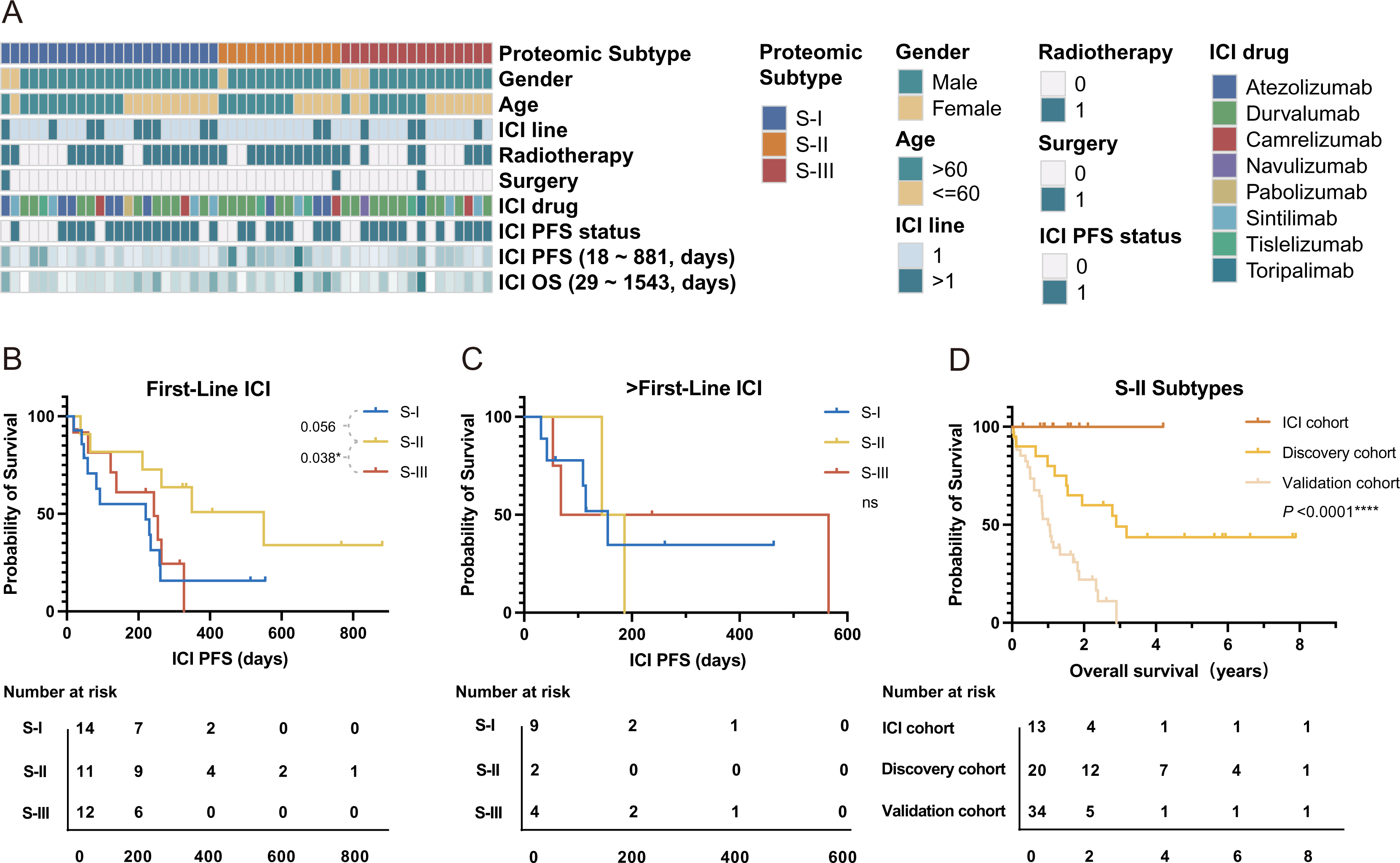
A. the clinical characteristics of the patients from the immunotherapy cohort. **B-C.** In the immunotherapy cohort, the S-II patients had the best PFS time when receiving first-line immunotherapy (**B**) but showed no statistically obvious difference between S-I/III when receiving second-line immunotherapy (**C**). **D.** In all S-II patients, patients had better overall survival after immunotherapy compared with those did not received immunotherapy.

### Potential drug repurposing targets for S-III subtype patients

Since patients in the S-III subtype had the worst OS and did not benefit from chemotherapy, they were the most in need of new therapeutic options. We selected specifically overexpressed proteins in S-III compared with S-I and S-II and investigated their feasibility as potential drug repurposing targets. We first identified previously investigated actionable drug targets and found that at least one target among EGFR, AURKB, BCL2, and EZH2 was highly expressed in a total of 21/27, or 77.8% of S-III patients. Drugs targeting AURKB, BCL2, or EZH2 are currently in various stages of clinical development and have shown promising results for treating certain cancers [34,35,36]. Many kinases, phosphatases, transporters, UPS proteins, and other enzymes were overexpressed in S-III, accounting for nearly 1/3 of all specifically overexpressed proteins in S-III. Proteins that regulate necroptosis, including MLKL, TRAF2, and RIPK1 are also potential targets [37]. Their druggability needs to be further investigated. An overview of the potential drug repurposing targets for each individual patient is shown in **Figure 5**.

**Figure 5.**
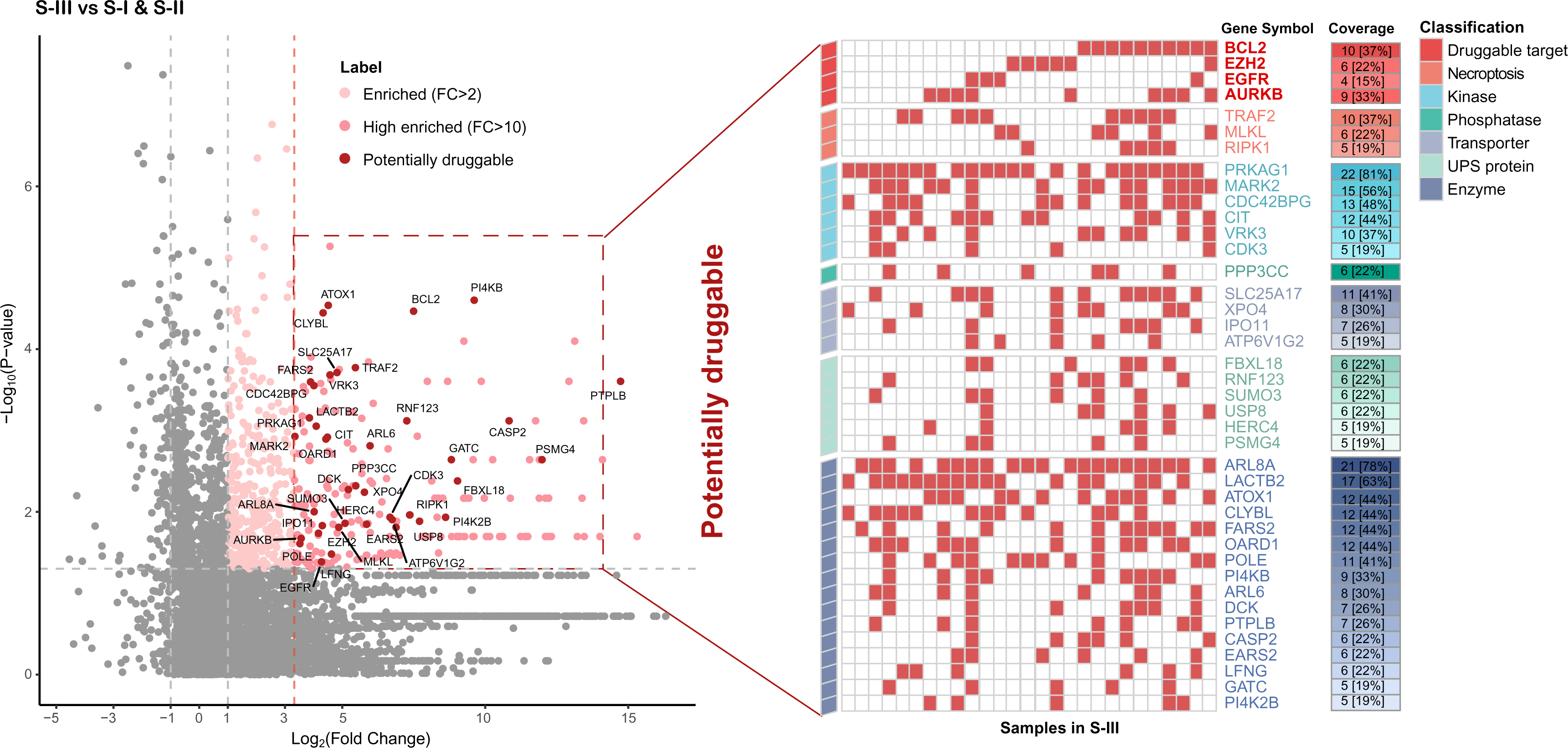
The atlas of potential drug targets for individual patients in S-III.

## Discussion

While genomic and transcriptomic analyses have greatly improved our understanding of SCLC tumorigenesis, their values in identifying effective stratification markers and therapeutic targets are limited. In this retrospective proteomic analysis, we showed that proteomics alone could stratify SCLC for prognosis and chemo-sensitivity. Since most SCLC tumors are non-resectable when diagnosed, we validated our proteomic subtyping using biopsy samples from an independent center, expanding the clinical utility of our proteomic subtyping method. Our analysis shows that all S-I patients in the discovery cohort who received chemotherapy were alive by the end of the study, confirming the suitability of chemotherapy as first line treatment option for these patients. Antigen presentation and other immune responses were enriched in S-II, suggesting that immune checkpoint inhibitors (ICIs) may be a viable choice for S-II patients. We validated this prediction in a third independent group of patients that received immunotherapy as the first-line treatment. Notably, a previous transcriptomic study also defined an “inflamed” SCLC subtype (SCLC-I) [15] characterized by high expression of genes related to HLAs and experienced greatest benefit from the addition of anti-PDL1 to chemotherapy [15]. Our proteomic subtyping thus may provide value in guiding SCLC treatment.

Since neither PDL1 expression nor TMB is a reliable biomarker for the prediction of immunotherapy response in clinical trials such as IMPower133 and CheckMate032, other biomarkers are in great need. Our analysis showed that patients in the S-II subtype may benefit from the immunotherapy. Although they were mostly ES-SCLC and lost the oppotunity for surgery, the two-year survival rate of the S-II patients with immunotherapy reached over 30%, much better than the other subtypes. In other clinical trials (for example, ASTRUM-005), the median PFS was generally less than 6 months [38], which was similar to that of S-I/III patients in our study. However, the median PFS reached around one year for the S-II subtypes here. It will be important to investigate in a prospective trial to determine whether the S-II subtype could serve as a new predictive biomarker for guiding SCLC immunotherapy.

Mechanisms of immune resistance vary [39]. MHC-I molecules were generally expressed at low levels in the SCLC tumor cells, resulting in low antigen presentation. In addition, tumor cells can also secrete factors to inhibit antigen presenting cells. About 50% of patients with SCLC have almost no T cell infiltration, and there are also suppressive immune cells in the immune microenvironment of SCLC. In our study, S-II subtype had both higher MHC-I molecules, along with other proteins in antigen presenting. Interferon gamma signaling was also enriched in S-II. Thus, S-II may be the immune hot subtype while S-I/III were the cold ones.

Among proteins and pathways as potential drug targets for the S-III subtype, BCL2, EZH2, ARUKB, and EGFR are actionable drug targets that have been at various stages of clinical trials. Among them, EZH2 was identified as an upstream regulator in the SLFN11 axis that mediates acquired chemoresistance in an *in vitro* PDX model [40]. FDA has recently approved the EZH2 inhibitor tazemetostat for treating epithelioid sarcoma [41]. Interestingly, EZH2 is a negative regulator of MHC class I molecules, and inhibiting EZH2 could enhance antigen presentation and circumvent anti-PD-1 resistance [42].

Although EGFR-TKi (such as Gefitinib) was not effective in a SCLC clinical trial [43], the identification of a subgroup of patients with EGFR overexpression suggest an alternative approach by using EGFR antibody. Notably, EGFR antibody has been approved as first-line treatment option for KRAS wildtype, EGFR overexpressing colon cancer patients [44]. BCL2 family proteins comprise the sentinel network that regulates mitochondrial and intrinsic apoptotic responses. Previous studies of a BCL-2 inhibitor fell short of expectations in SCLC clinical trials [45]. A possible explanation is that patients in the trial were not screened for BCL-2 expression. Interestingly, the SCLC-A transcriptomic subtype [15], which had higher expression of BCL2, was found to be sensitive to multiple BCL2 inhibitors in an *in vitro* study. Recently, the single target BCL2 inhibitor Venetoclax, which was approved by the FDA for acute myeloid leukemia and chronic lymphocytic leukemia [46], showed therapeutic effect on multiple patient-derived xenografts of SCLC [47]. AURKB is a mitotic protein kinase. Phase I/II clinical studies of its inhibitor Alisertib have demonstrated increased antitumor effects in various hematologic malignancies and solid tumors [48].

In addition, our proteomic data suggested that necroptosis regulators including MLKL, TRAF2, and RIPK1 may be investigated as potential therapeutic targets for SCLC. Necroptosis causes cellular swelling and plasma membrane collapse, which may lead to the release of intracellular biomolecules including damage-associated molecular patterns (DAMPs) and cytokines. These molecules can perform immunological functions such as chemotaxis, phagocytosis, and immune cell activation [49]. Although initial attraction of antigen presentation cells such as macrophages and dendritic cells (DCs) by DAMPs/cytokines could recruit CD8/CD4 positive T cells for immune activation in early stage, the recruitment of myeloid-derived suppressor cells (MDSC) and tumor-associated macrophages (TAM) at later stages could lead to immune suppression [50]. The effects of necroptosis proteins on tumors are cancer dependent. Cytokines released by necroptotic cancer cells can promote tumor angiogenesis, proliferation, and metastasis. For instance, high expression of RIPK1 was linked to metastasis in breast cancer [51] and poor survival in glioblastoma [52] but was a good prognostic indicator in head and neck cancer [53]. High MLKL expression correlated positively with good prognosis in colorectal cancer [54], HR-HPV cervical cancer (high risk-human papillomavirus) [55], ovarian cancer [56], and pancreatic adenocarcinoma [57], but negatively in breast cancer [58], cervical squamous cell carcinoma [59], and gastric cancer [60]. We found in this study a correlation between high expression of MLKL, TRAF2, and RIPK1 and the most malignant SCLC S-III subtype, suggesting that these proteins may represent good targets for treating SCLC.

The S-III subtype is also enriched in proteins involved in chromatin organization and transcriptional regulation. For example, CHAF1A, TERF2, and TOX4 can mediate chromatin assembly, protect chromosome ends, and regulate chromatin binding during DNA replication [61], metaphase [62], and transition to interphase [63], respectively. For transcriptional regulation, ATAD2 and ZNF516 act as transcription activators, promoting the expression of CCND1/MYC/ E2F1 [64] or genes related to cellular response to replication stress [65], respectively. Moreover, PHF8 acts as a coactivator of rDNA transcription by activating polymerase I (pol I) mediated transcription of rRNA genes [66], while MED6 is a coactivator involved in the regulated transcription of nearly all RNA polymerase II-dependent genes [67]. These findings underline the importance of investigating dysfunctional chromatin organization in the development and treatment of SCLC.

Our study has some limitations. For example, our discovery cohort was relatively small compared with other omics studies, such as those of NSCLC. This is in part because most SCLC cases were detected at late stages without opportunity for surgery, so large-scale proteomic analysis of surgically resected samples was rare in the past. Moreover, our proteomic stratification was different from previous transcriptional classification based on ASCL1, NeuroD1, and YAP1, thus providing limited mechanistic insight into the tumorigenesis. The reason for the discrepancy is unclear, but we did notice by IHC the overlap of several of these markers.

In summary, our results demonstrate the validity and importance of using both surgical and biopsy FFPE samples and serve as a valuable resource for the SCLC research community. This work should also help guide future pre-clinical investigations that seek more meaningful and efficacious stratification of SCLC in order to improve therapeutic responses and patient survival.

## Materials and methods

### Sample collection

Tumors in the discovery cohort (diagnosed from 2012 to 2018) were obtained with informed consent from archival sources at Tongji Hospital, Tongji Medical College, Huazhong University of Science and Technology. Tumors in the second cohort (diagnosed from 2018 to 2021) were biopsy samples obtained from the First Affiliated Hospital of Henan University. The 52 samples in the immunotherapy cohort was also from Tongji Hospital (diagnosed from 2017 to 2021). The samples were collected from patients at initial diagnosis. All diagnoses were independently reviewed by three experienced pathologists, and complied with the latest World Health Organization classification standards. The tumors came from patients not treated with neoadjuvant chemotherapy or radiotherapy before operation, with no previous history of malignancy, having SCLC as the initial primary cancer diagnosis at the time of surgical resection, and with adequate tumor/tissue material as well as clinical annotation and follow-up time. The tumor cell content were >90% tumor cells judged by haematoxylin and eosin (H&E) staining and yielded >700 proteins in MS analysis. All cases were staged according to the National Comprehensive Cancer Network (NCCN) clinical practice guidelines in oncology for small cell lung cancer (version 4.2020).

### Immunohistochemistry and assesment

Immunohistochemical staining was conducted using tissue arrays. Prior to de-paraffinization, the slides were heated to 60°C for ten minutes to melt the paraffin. The slides were then washed three times with xylene to solubilize and remove the paraffin. Next, the xylene was removed by washing three times with 100% ethanol followed by 75%, 50% ethanol, and PBS. After the sections were deparaffinized and hydrated, the endogenous peroxidase activity was blocked. Antigen retrieval was performed using the Dako Target Retrieval Solution, High pH (Dako Ominis, Agilent Technologies, CA, USA), in a PTLink set at 98°C for 25 min. The slides were then incubated with the primary antibody at 4 ℃ overnight.

Primary antibodies used in this study were: anti-ASCL1 antibody (Abcam, ab211327, 1:200, Boston, MA), anti-NeuroD1 antibody (Abcam, ab213725, 1:200,), and anti-YAP1 antibody (Abcam, ab52771, 1:50,). Following secondary antibody incubation (∼1 hour), the sections were visualized with the DAB kit (ZLI-9017, Zhongshan Biotechnology, China) and counterstained with hematoxylin. The stained samples were scored by three pathologists independently for the multiplication of staining intensity (1-weak; 2-moderate; 3-strong) and the percentage of positive tumor cells, which resulted in scores of 0-300. A score of < 10 was designated as 0, 10-40 as 1+, 41-140 as 2+, and 141-300 as 3+. All samples with scores of > 10 were considered positive cases.

### Protein extraction, trypsin digestion, and LC-MS/MS processing

For each SCLC sample, proteins were extracted from three tissue slices (5-µm thick) from the FFPE block (2-5 mm in diameter) The de-paraffinization procedure was the same as IHC as described above. Air dried sample was scraped from the slide and resolubilized in 100 ml of 50 mM NH_4_HCO_3_. The sample was then incubated at 95 °C for 5 min for de-crosslinking, and cooled to room temperature. Trypsin digestion was carried out at 37°C overnight. Peptides were extracted twice with 200 μl of extraction buffer (50% acetonitrile and 0.1% formic acid in water) with 15 min vortex. The resulting peptides were dried and stored at -80 ° C for future analysis. The digested peptides were eluted, divided into three fractions, and analyzed on a Q Exactive HF-X or Orbitrap Exploris™ 480 mass spectrometer coupled with an Ultimate 3000 RSLCnano LC system (Thermo Fisher Scientific) and operated at data-dependent aquisition mode. MS1 was measured in the Orbitrap at a resolution of 60,000 followed by tandem MS scans of the top 40 precursors using higher-energy collision dissociation with 27% of normalized collision energy and 15 s of dynamic exclusion time.

### Mass spectrometry data analysis

MS Raw files were searched against the National Center for Biotechnology Information (NCBI) Ref-seq human proteome database (updated on 04/07/2013, 27,414 entries) in Firmiana [68], a one-stop proteomic cloud platform for data processing and analysis, implemented with Mascot search engine with Percolator (Matrix Science, version 2.3.01). The following search parameters were used: (1) Mass tolerances were 20 ppm for precursor ions and 0.05 Da for product-ions; (2) Up to two missed cleavages were allowed; (3) The minimal peptide length was seven amino acids; (4) Cysteine carbamidomethylation was set as a fixed modification, and N-acetylation and methionine oxidation were considered variable modifications; and (5) The charges of precursor ions were limited to + 2, + 3, + 4, + 5, and + 6. The peptide and protein FDR were both set to 1%. A label-free, intensity-based absolute quantification (iBAQ) algorithm was used for protein quantification. The iBAQ values were calculated by dividing the raw intensities by the number of theoretical observable peptides. FOT (fraction of total), calculated by dividing a protein’s iBAQ by the sum of iBAQs of all identified proteins in a single experiment, was used as normalized abundance to compare protein abundance across all experiments. The missing value was imputed with 1/10 of the global non-zero minimum value of the sample [69].

### Subtyping and validation

A non-negative matrix factorization (NMF) consensus clustering algorithm was used for subtyping the discovery dataset. The standard “brunet” option was selected and 50 iterations were performed. The number of clusters (k) was set as 2 to 6, and the minimum member of each subclass was set as 10. The silhouette indicator (Average silhouette > 0.8) and prognostic association (*P*-value < 0.01) were used to determine the optimal clustering number.

To develop a robust classification model, we first selected the top 1,100 most abundant proteins detected from each sample, which yielded a dataset of 3,460 proteins; then the 445 proteins that were detected in at least 7 samples (>10%) with the coefficient of variations (CV) greater than 1.9 were used for non-negative matrix factorization (NMF) consensus clustering (Table S3) To validate the classification in an independent external dataset, the random forest (RF) classifier was implemented on the discovery set using the R function randomForest (package: randomForest). The top 500 most abundant proteins detected in each sample were aggregated for differential expression analysis. Finally, 58 significantly differentially expressed proteins (Fold change > 1.5, adjusted t-test *P*-value < 0.05) with high identification frequencies (detected in more than 25% of the samples) were used as input (predictor variables). The subtype terms of SCLC on the discovery set were used as the response variables. The optimal parameters were estimated from the R function train (package: caret). The 10-fold cross-validation strategy was utilized for internal validation.

### Survival analysis

The Kaplan-Meier method, log-rank test, and the Cox proportional-hazards model with Wald statistics were used for survival analysis in all datasets. The multivariate Cox analysis was adjusted for age, sex, staging system, and chemotherapy. Hazard ratios (HRs) with 95% confidence intervals (CI) were estimated for each variable. All calculated *P*-values were two-sided where <0.05 was considered statistically significant.

### Bioinformatic analysis

To investigate the proteomic features of SCLC subtypes, we selected significantly altered proteins by comparing their expression in each subtype to the other subtypes (Wilcox test *P* < 0.05 and fold change > 3 for S-I and II or fold change > 10 for S-III). Metascape analysis was conducted online (https://metascape.org) for functional pathway analysis. The on-line tool (Sangerbox tools, http://www.sangerbox.com/tool) was used for generating the plots and heatmap. All other statistical analyses were carried out with R 3.6.1.

### Ethical statement

This study was approved by the ethical committee of Tongji Hospital, Tongji Medical College, Huazhong University of Science and Technology (Approval Number: TJ-IRB20201011).

#### Data availability

All data and search results have been deposited to the iProX database (http://www.iprox.org) with the iProX accession: IPX0004230000.

## Acknowledgments

This work was supported by the National Key R&D Program of China (Grant Nos. 2018YFA0507503, 2017YFA0505102, 2017YFA0505103, and 2017YFA0505104), the National Natural Science Foundation of China (Grant Nos. 82072597, 62131009, 31770892, 31970725, 31870828, 81874237, and 81974016), Beijing Municipal Natural Science Foundation (Grant No. 7192199), and State Key Laboratory of Proteomics (Grant No. SKLP-K202002), Kaifeng science and technology development plan project (Grant No. 1806005). We thank the Mass Spectrometry Platform of National Center for Protein Sciences (Beijing) (Phoenix Center) for technical assistance. We also thank all the patients who participated in this study, as well as the clinicians and staff of the institutions who assisted with the studies.

## Declaration of AI and AI-assisted technologies

No AI and AI-assisted technologies were used in writing this paper.

### CRediT author statement

**Zitian Huo:**Writing - original draft,Formal analysis, Investigation,Visualization

**Yaqi Duan:**Writing - review & editing.Methodology.

**Dongdong Zhan:**Investigation, Data curation,Validation

**Xizhen Xu:**Visualization,Writing - review & editing.

**Nairen Zheng:**Data curation. Validation.

**Jing Cai:**Data curation. Resources.

**Ruifang Sun:**Resources, Data curation.

**Jianping Wang:**Resources, Data curation.

**Fang Cheng:**Resources, Data curation.

**Zhan Gao:**Resources, Data curation.

**Caixia Xu:**Resources, Data curation.

**Wanlin Liu:**Resources, Data curation.

**Yuting Dong:**Resources, Data curation.

**Sailong Ma:**Resources

**Qian Zhang:**Resources

**Yiyun Zheng:**Resources

**Liping Lou:**Resources

**Dong Kuang:**Writing - review & editing.

**Qian Chu:** Supervision, Project administration, Funding acquisition.

**Jun Qin:**Conceptualization, Resources, Supervision, Project administration, Funding acquisition.Writing - review & editing.

**Guoping Wang:**Conceptualization, Resources, Supervision, Project administration, Funding acquisition.

**Yi Wang:**Conceptualization, Resources, Supervision, Project administration, Funding acquisition.Writing - review & editing.

All authors read and approved the final manuscript.

## Supplementary material

**Figure S1.**
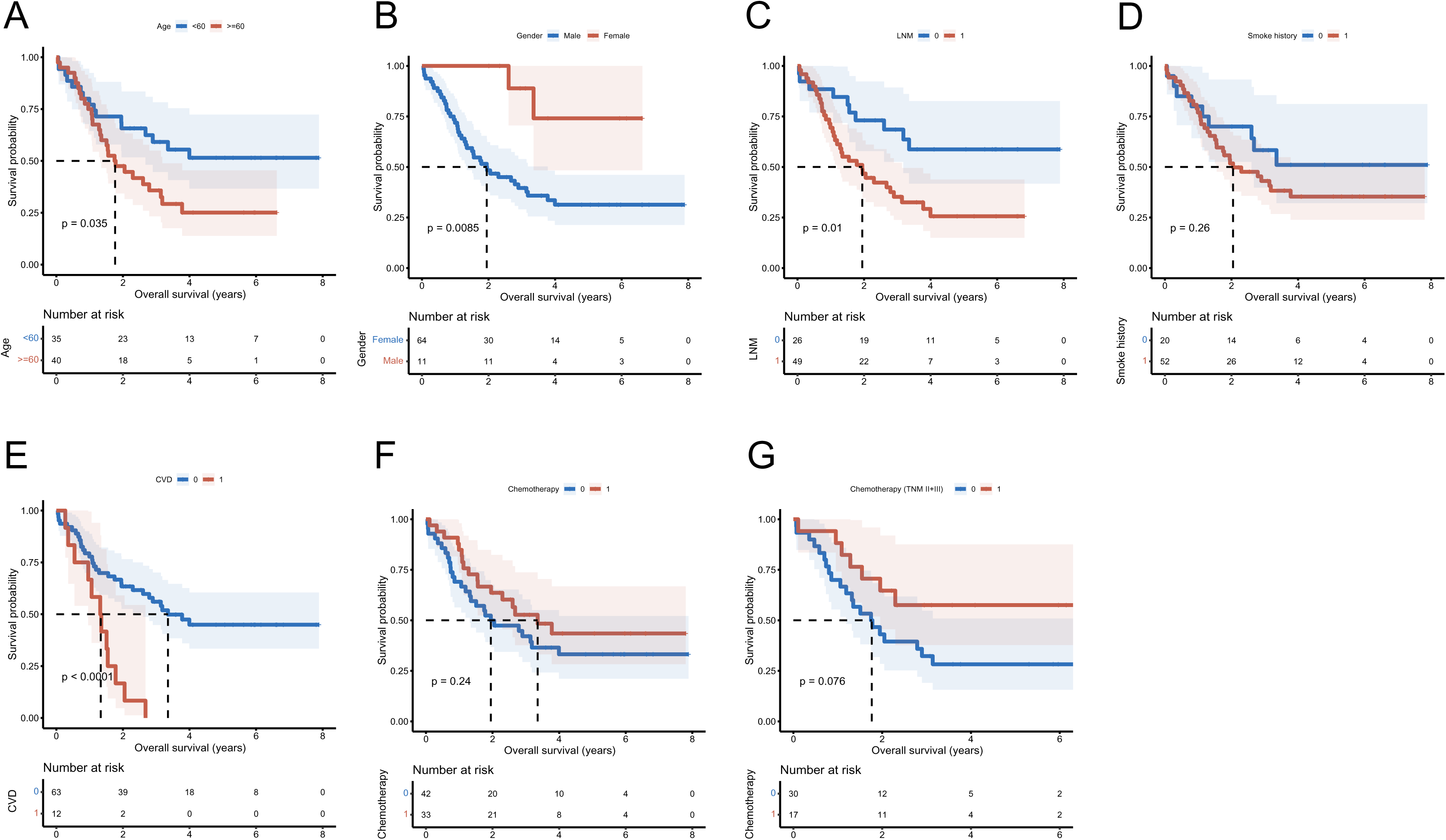
Univariate prognostic analysis of common clinical pathological factors.

**Figure S2.**
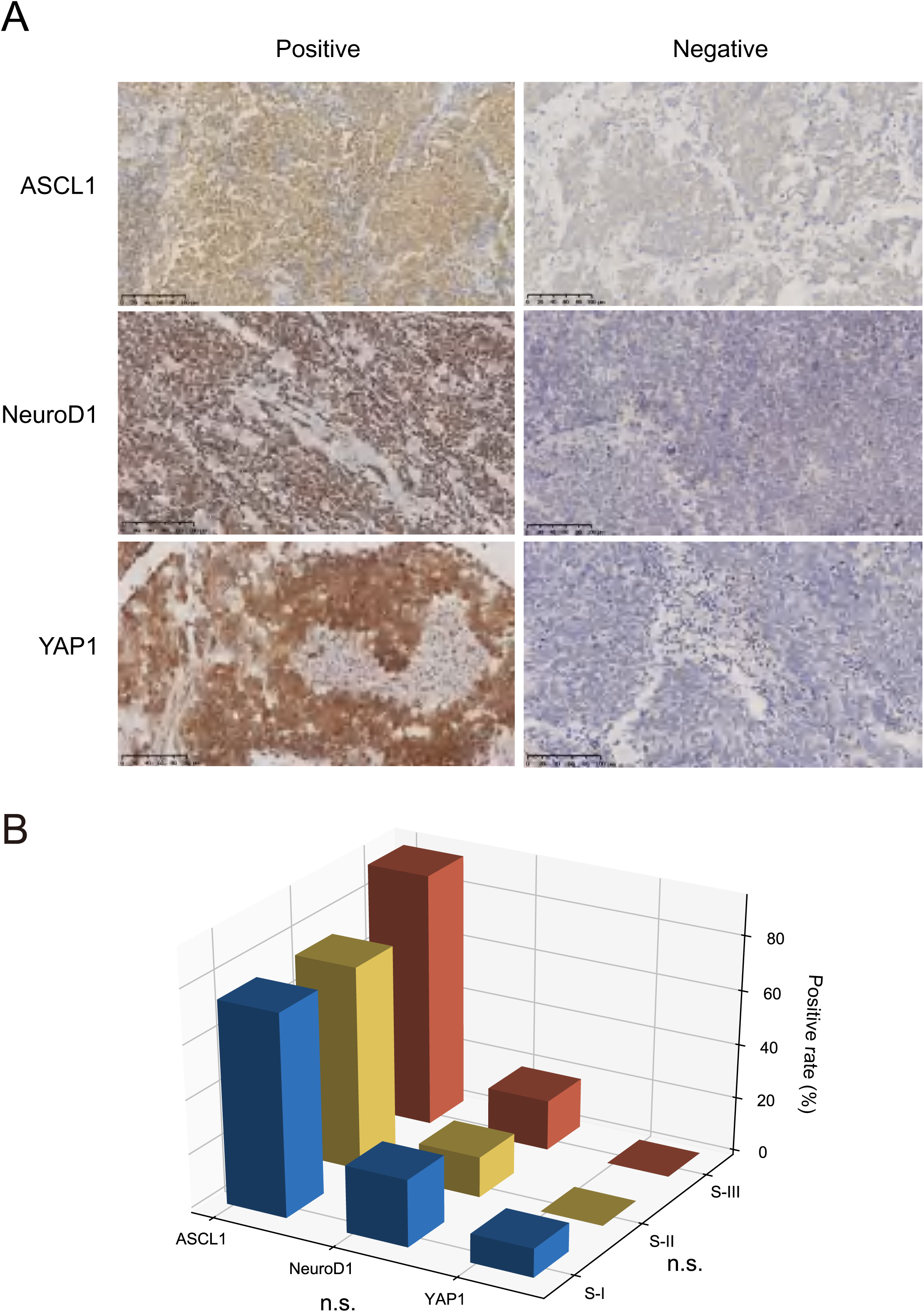
Representative IHC staining and distribution of common neuroendocrine biomarkers in SCLC. **A.** Representative IHC staining of ASCL1, NeuroD1, and YAP1. B. The expression of ASCL1, NeuroD1, and YAP1 was not distributed differently across the three subtypes.

**Figure S3.**
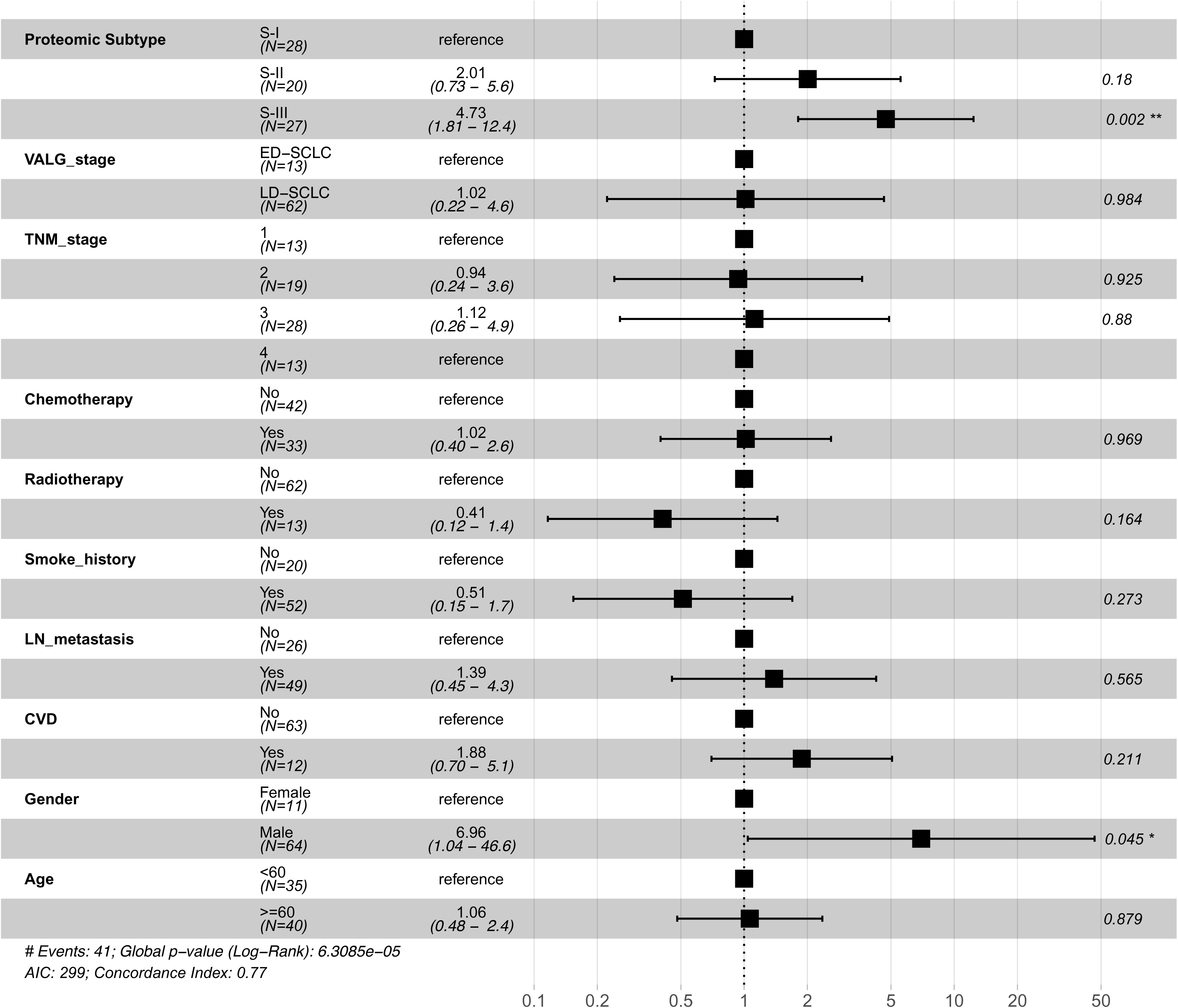
**SCLC** proteomic subtype was an independent prognostic factor confirmed by multivariate Cox analysis.

**Table S1.** Clinical- pathological data of discovery cohort.

**Table S2.** Proteomic results(FOT) of each case in the discovery cohort.

**Table S3.** Proteins for the non-negative matrix factorization (NMF) consensus clustering.

**Table S4.** Fifty-eight signature proteins for the predictive classifier model.

**Table S5.** Clinical- pathological data of validation cohort.

**Table S6.** Proteomic results(FOT) of each case in the validation cohort.

**Table S7.** Signature proteins for each subtype.

**Table S8.** Clinical- pathological data of the immunotherapy cohort.

**Table S9.** Proteomic results(FOT) of each case in the immunotherapy cohort.

